# Sepsis: Partial least squares structural equation modelling (PLS-SEM) suggests a critical role for anti-inflammatory responses in clinical severity

**DOI:** 10.1101/193615

**Authors:** Ratnadeep Mukherjee, Pijus Kanti Barman, Pravat Thatoi, Birendra Kumar Prusty, Rina Tripathy, Bidyut Kumar Das, Balachandran Ravindran

**Author notes:** Corresponding author: Balachandran Ravindran, Director, Institute of Life Sciences, NALCO Square, Chandrasekharpur, Bhubaneswar–751023, Odisha, India.

## Abstract

**Backgorund:** Despite major advances in medicine, Sepsis remains one of the major killers in critical care wards around the world. For several years it was widely believed that an early pro-inflammatory host response is followed by an overwhelming anti-inflammatory phase. The hypo-inflammatory status, termed as ‘Compensatory anti-inflammatory response syndrome’ (CARS), was proposed to be the primary cause of sepsis-associated mortality. However, this paradiam changed in recent years since there was little evidence to support the linear model of host response and pathogenesis in sepsis. Currently held view is that both inflammatory and anti-inflammatory host responses are stimulated in an overlapping manner. In this study a robust statistical model to study the complex interplay of host cytokines in human sepsis has been developed to evaluate host responses in sepsis that contribute significantly to clinical pathology.

**Methods:** Twentyseven cytokines/ chemokines were quantified in 139 sepsis patients and multivariate analysis of variance (MANOVA) was performed to assess differences in host responses in different categories of clinical severity. Partial least squares regression based structural equation modelling (PLS-SEM) was used to assess interactions between different groups of cytokines and their contribution to clinical pathology. An array of 23 cytokines was analysed in a mouse model of endotoxemia and a similar mathematical model was constructed.

**Results:** The results of MANOVA demonstrated the ability of combined cytokine response to discriminate sepsis patients according to clinical severity or outcome. Structural equation modelling revealed strong positive association between inflammatory and anti-inflammatory cytokines. In human sepsis, anti-inflammatory cytokines emerged as a significant entity associated with clinical severity as assessed by APACHE II scores.

**Conclusion:** PLS-SEM modeling of cytokine responses and APACHE II score in human sepsis revealed that anti-inflammatory molecules contribute significantly towards clinical severity. More critically, the model offers emperical evidence for failures of clinical trials conducted during the last two decades in which antagonists of inflammatory host responses for human Sepsis were used for sepsis. The model also provides credence to the notion that inflammatory and anti-inflammatory host responses occur concurrently in both experimental endotoxemia and in human sepsis.

## Background

Sepsis is a complex clinical syndrome associated consistently with an unregulated and exaggerated inflammatory response to infection. It remains a major health problem with an overall hospitality rate of about 30%, which has remained nearly stable during the last two decades [1]. For many years, the primary reason for sepsis pathogenesis was thought to be hyperinflammation that led to numerous attempts in the last three decades focussing on molecules that would block hyperinflammation. These have included antagonists to TLR4, CD14, RAGE, TNF-α, iNOS and inflammasome activation [2-6]. However, all such attempts, although promising initially in experimental animal models failed in clinical trials undertaken in patients [7].

Failure of management of human sepsis by decreasing hyper-inflammation has resulted in a paradigm shift in the way immune response to sepsis is perceived currently. Host response to sepsis was proposed to be biphasic consisting of an initial systemic inflammatory response syndrome (SIRS) characterized by excessive production of a large number of pro-inflammatory mediators followed by a compensatory anti-inflammatory response syndrome (CARS), which involves excessive release of immunosuppressive mediators, de-activation of monocytes, T cell anergy, and loss of immune cells by apoptosis [reviewed in 8]. While overwhelming inflammation may cause organ injury and shock, prolonged hypo-inflammatory state may lead to immunosuppression and failure to clear secondary infections [9-11]. This has led to the notion that immunosuppression due to sepsis could be more critical for adverse prognosis (than the initial hyperinflammatory phase) by rendering the patients susceptible to otherwise weakly virulent pathogens [12]. In spite of these insights, novel therapies targeting this aspect of sepsis pathogenesis were not attempted vigouraouly due to concerns that such therapies could aggrevate detrimental effects of hyperinflammation [13]. However, in recent times, intervention strategies using immunostimulatory cytokines have gained increasing acceptance [12]. One such cytokine is IL-7, which showed promise in a preclinical experiment demonstrating improved lymphocyte proliferation, IFN-γ production and Bcl-2 induction of patient cells treated ex vivo with IL-7 [14]. In another study, IL-15 treatment was shown to decrease immune cell apoptosis by increasing Bcl-2 expression in septic mice, thereby making it yet another novel immunomodulatory cytokine of interest [15]. The third important cytokine that demostrated benefits in anit-septic therapy is GM-CSF. Two recent Phase II clinical trials involving GM-CSF administration in patients with low HLA-DR expression on monocytes revealed improved illness score, enhanced blood monocyte HLA-DR expression and improved cytokine production [16].

While existing literature does not offer credence to the view of bimodal occurrence of SIRS and CARS during the course of sepsis, there are reports demonstrating concurrent induction of inflammatory and immunosuppressive molecules in sepsis [17, 18]. This has led to coining of a new expression, mixed antagonist response syndrome (MARS), a state in which both pro-and anti-inflammatory phenotypes coexist. However, role of MARS in determining disease severity and outcome in human sepsis is still unclear. In this study we have attempted multivariate statistical modelling of high-dimensional plasma cytokine data that demonstrates simultaneous induction and regulation of inflammatory and anti-inflammatory responses during sepsis. The analysis and statistical modelling suggests that the compensatory anti-inflammatory response rather than systemic inflammation is more critically responsible for pathogenesis and adverse prognosis in sepsis.

## Methods

### Reagents and kits

Gram-negative bacterial lipopolysaccharide (LPS, E.coli O55:B5) was obtained from Sigma chemicals. Acetic acid, citric acid, and dextrose were all from Fisher Scientific. Human 27-plex cytokine and Mouse 23-plex cytokine analysis kits were procured from Bio-Rad Laboratories.

### Human sepsis patients

The study was approved by ethics committees of Institute of Life Sciences and S.C.B. Medical College, and signed informed consent was obtained from all participants. 139 sepsis patients who were admitted to medical intensive care units at the Department of Medicine, S.C.B. Medical College and Hospital (Cuttack, India) were recruited for the study. For sepsis, patients were eligible for inclusion only if they had systemic inflammatory response syndrome and had a source of infection, proven or suspected. The acute physiological and chronic health evaluation II (APACHE II) scoring system was used to categorize the patients. Definitions of sepsis, severe sepsis, septic shock, and multi-organ dysfunction syndrome (MODS) were in accordance with published criteria [19, 20]. The following categories of patients were excluded from the study: patients with diabetes mellitus, hypertension, nephrotic syndrome, chronic kidney disease (sonographic feature of CKD and/or GFR<30 ml/min), patients with cardiac failure and immunocompromised individuals. Blood was collected in vials containing 15% (v/v) Acetate Citrate Dextrose (ACD). This was followed by isolation of plasma by centrifugation at 2000 rpm for 10 min. Plasma was stored in single-use aliquots at − 80°C.

### Mouse model of endotoxemia

8-10 weeks old male C57BL/6 mice were used for the study. All animal experiments were approved by the institutional animal ethics committee of Institute of Life sciences, Bhubaneswar. To simulate two groups based on lethality in human sepsis cases, a 5mg/kg nonlethal dose and 35mg/kg lethal dose of gram negative bacterial lipopolysaccharide (E.coli O55:B5, Sigma) was injected intraperitoneally. The animals were sacrificed at 2, 4, 8 and 16 hours post injection to mimic early and late stages of endotoxemia. Blood was collected in vials containing ACD as anticoagulant (15% v/v). Collected blood was centrifuged at 2000 rpm for 10 minutes for isolation of plasma. Isolated plasma was stored in 100 μl single-use aliquots at −80°C.

### Measurement of cytokines in plasma

Human plasma was analysed using the human 27-plex cytokine panel (Bio-Rad) according to manufacturer’s instructions and contained the following targets: IL-1β, IL-1ra, IL-2, IL-4, IL-5, IL-6, IL-7, IL-8, IL-9, IL-10, IL-12(p70), IL-13, IL-15, IL-17, Basic FGF, Eotaxin, G-CSF, GM-CSF, IFN-γ, IP-10, MCP-1, MIP-1α, MIP-1β, PDGF, RANTES, TNF-α, and VEGF. Mouse plasma was analysed using the mouse 23-plex cytokine panel (Bio-Rad) as specified by the manufacturer and contained the following targets: IL-1α, IL-1β, IL-2, IL-3, IL-4, IL-5, IL-6, IL-9, IL-10, IL-12(p40), IL-12(p70), IL-13, IL-17, Eotaxin, G-CSF, GM-CSF, IFN-γ, KC, MCP-1, MIP-1α, MIP-1β, RANTES, and TNF-α. All samples were read on a Bioplex 200 system (Bio-Rad). Concentrations of unknown samples were interpolated from an 8-point standard curve fitted with a five-parameter logistic regression model.

### Descriptive statistics and exploratory data analysis

For comparison between two groups, a nonparametric Man – Whitney U –test was conducted. One – way analysis of variance (ANOVA) followed by Bonferroni’s multiple comparison test was carried out to compare means between three or more groups. Exploratory data analysis (EDA) methods were based on construction of a n X n - dimensional correlation matrix to assess associations between measured response variables. Association of plasma cytokines with morbidity and mortality in sepsis was investigated using multivariate analysis of variance (MANOVA). The Wilk’s lambda statistic was used to test significance of among-group differences, while post hoc procedures based on pair-wise contrasts and canonical analyses were used to respectively determine which of the various groups actually differed from each other and which responses and what type of relationships among them were responsible for the observed inter-group differences. All response data were logarithmically transformed {loge (*n*+1)} prior to performing multivariate statistical analyses to satisfy linearity assumption. All statistical analyses were performed in R programming environment for statistical computing. For performing canonical analysis, package candisc was used. For all statistical comparisons, a value <0.05 was considered statistically significant.

### Structural Equation Modelling

To predict which immune response variables are most responsible for clinical pathology in endotoxemia or sepsis, partial least squares (PLS) regression – based structural equation model (PLS-SEM) was constructed taking cytokines as predictors and APACHE II score as outcome. The measured cytokine concnetrations in plasma were loge(n+1) transformed and were grouped into the following latent variables based on existing knowledge: IL-1β, TNF-α, IL-6 and IL-12 for SIRS; IL-4, IL-10 and IL-13 for CARS; IFN-γ, MIP-1α and MIP-1β for Th1 response; IL-4, IL-13; Eotaxin for Th2 response; IL-2, IL-4, IL-5, IL-7; IL-9 for T and B cell proliferation and IL-5 and GM-CSF for Hematopoiesis. Cytokines with factor loadings < 0.5 were excluded from the model. For assessing significance of strength of interaction between each predictor and outcome variable pair, bootstrapping was performed. Bootstrapping values in excess of 1.9 were considered statistically significant at p<0.05. PLS-SEM analysis was conducted in SmartPLS v3.0 software.

## Results

### Patient characteristics and descriptive statistics

139 patients were included in the study group (95 men and 44 women). Table 1 summarizes the demographic and clinical characteristics. About 35% (n = 48) individuals were deceased out of the total cohort. More than 60% of the cases (66 out of 139) were diagnosed with severe sepsis at the time of admission. Incidence of sepsis and septic shock were almost equally distributed in the cohort (28% and 25% respectively).

**Table 1.**
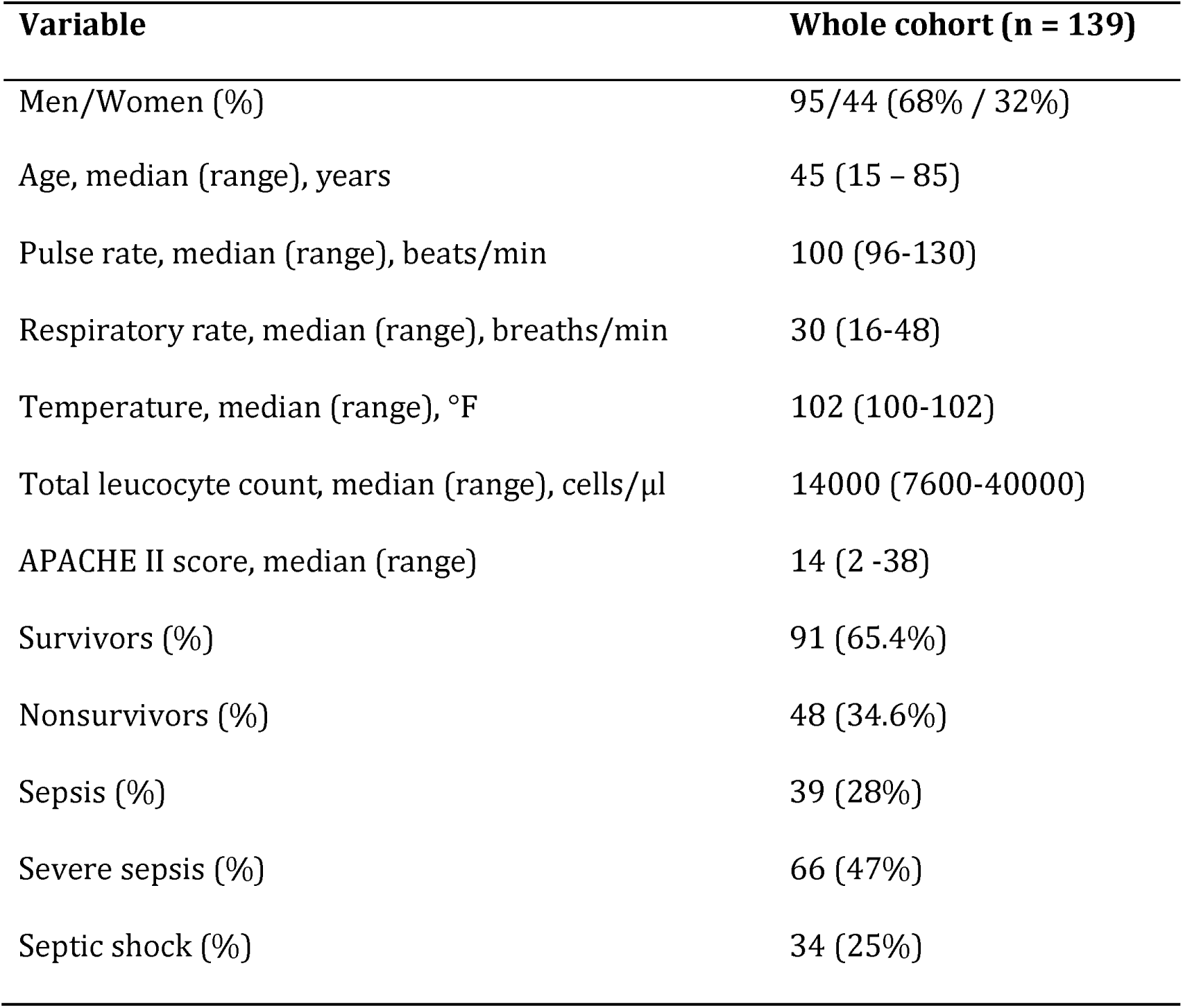
Demographic and clinical characteristics of patients

| **Variable** | **Whole cohort (n = 139)** |
| --- | --- |
| Men/Women (%) | 95/44 (68% / 32%) |
| Age, median (range), years | 45 (15 – 85) |
| Pulse rate, median (range), beats/min | 100 (96-130) |
| Respiratory rate, median (range), breaths/min | 30 (16-48) |
| Temperature, median (range), °F | 102 (100-102) |
| Total leucocyte count, median (range), cells/μl | 14000 (7600-40000) |
| APACHE II score, median (range) | 14 (2 -38) |
| Survivors (%) | 91 (65.4%) |
| Nonsurvivors (%) | 48 (34.6%) |
| Sepsis (%) | 39 (28%) |
| Severe sepsis (%) | 66 (47%) |
| Septic shock (%) | 34 (25%) |

Summary statistics of all individual cytokines discriminated based either on outcome or clinical category is presented in Tables 2 and 3 respectively. Of all the measured cytokines, IL-1β, IL-2, IL-4, IL-5, IL-17 and IFN-γ were able to discriminate between survivors and nonsurvivors and also between clinical categories of sepsis. On the other hand, plasma levels of BasicFGF and RANTES could only differentiate survivors from nonsurvivors whereas IL-7, IL-9, IL-15, Eotaxin, GM-CSF, MIP-1α, MIP-1β and TNF-α levels were successful only in discriminating different clinical stages of sepsis. All cytokines that were able to segregate survivors and non-survivors showed significantly elevated levels in plasma of survivors suggesting a possible immune response deficiency among non-survivors. Similarly, the cytokines that were able to contrast clinical categories of sepsis showed elevated levels in patients with sepsis as compared to severe sepsis and septic shock. These two findings are in agreement with each other as number of deaths in patients with severe sepsis and septic shock (32 out of 66 and 13 out of 34 respectively) far outnumbered cases of death in sepsis category (3 out of 39).

**Table 2.** Comparison of plasma cytokine levels in survivors and nonsurvivors

|  |  |  |  |
| --- | --- | --- | --- |
| **Cytokine concentration in pg/ml; median(range)** |  |  |  |
| **Cytokines** | **Survivors (n=91)** | **Nonsurvivors (n=48)** | **p-values** |
| IL-1β | 9.67 (0.00 - 3008.43) | 6.34 (0.00 - 4367.19) | **0.03774** |
| IL-1RA | 245.54 (0.00 - 2381.50) | 217.23 (4.54 - 1056.58) | 0.186 |
| IL-2 | 7.88 (0.00 - 98.93) | 3.74 (0.00 - 43.80) | **0.001273** |
| IL-4 | 7.10 (0.00 - 24.00) | 5.85 (0.00 - 23.00) | **0.0272** |
| IL-5 | 4.76 (0.00 - 87.60) | 1.24 (0.00 - 39.14) | **0.02272** |
| IL-6 | 91.77 (18.50 - 5952.67) | 112.13 (17.92 - 7260.00) | 0.4474 |
| IL-7 | 28.55 (0.00 - 615.45) | 17.91 (0.00 - 290.19) | 0.09795 |
| IL-8 | 199.92 (0.00 - 4078.00) | 187.25 (0.00 - 6681.00) | 0.9312 |
| IL-9 | 53.34 (0.00 - 202.00) | 44.66 (0.00 - 197.90) | 0.1049 |
| IL-10 | 21.96 (1.62 - 505.90) | 19.92 (1.59 - 149.62) | 0.4024 |
| IL-12 (p70) | 84.51 (8.80 - 2022.38) | 58.16 (0.00 - 278.01) | 0.05648 |
| IL-13 | 9.53 (0.00 - 238.96) | 8.66 (0.00 - 133.33) | 0.2642 |
| IL-15 | 12.98 (0.00 - 36.12) | 11.64 (0.00 - 97.40) | 0.8838 |
| IL-17 | 168.94 (0.00 - 579.91) | 119.95 (0.00 - 480.70) | **0.001886** |
| Eotaxin | 191.23 (26.76 - 728.16) | 158.09 (35.98 - 765.14) | 0.1493 |
| Basic FGF | 111.86 (29.08 - 7937.00) | 82.39 (0.00 - 7041.00) | **0.001788** |
| G-CSF | 105.35 (0.00 - 15425.17) | 84.66 (0.00 - 1504.81) | 0.05422 |
| GM-CSF | 45.10 (0.44 - 225.58) | 40.13 (0.44 - 237.64) | 0.2821 |
| IFN-γ | 527.58 (33.24 - 3120.00) | 356.14 (0.00 - 2335.70) | **0.04359** |
| IP-10 | 3681.07 (413.00 - 106944.21) | 2802.23 (90.35 - 173251.96) | 0.06968 |
| MCP-1 | 88.47 (23.52 - 1069.57) | 95.56 (0.00 - 4235.38) | 0.2758 |
| MIP-1α | 15.10 (0.00 - 2327.00) | 10.25 (0.00 - 2796.00) | 0.3276 |
| MIP-1β | 386.63 (0.00 - 10258.00) | 317.94 (0.00 - 6107.00) | 0.7197 |
| PDGF-BB | 2631.40 (52.39 - 14516.59) | 1271.10 (0.00 - 76944.00) | 0.07495 |
| RANTES | 6580.20 (257.87 - 1726538.00) | 4351.62 (0.00 - 1726537.60) | **0.02597** |
| TNF-α | 155.99 (14.57 - 10385.00) | 105.24 (0.00 - 4401.00) | 0.09101 |
| VEGF | 184.10 (20.59 - 1698.20) | 131.58 (0.00 - 2031.87) | 0.127 |

**Table 3.**
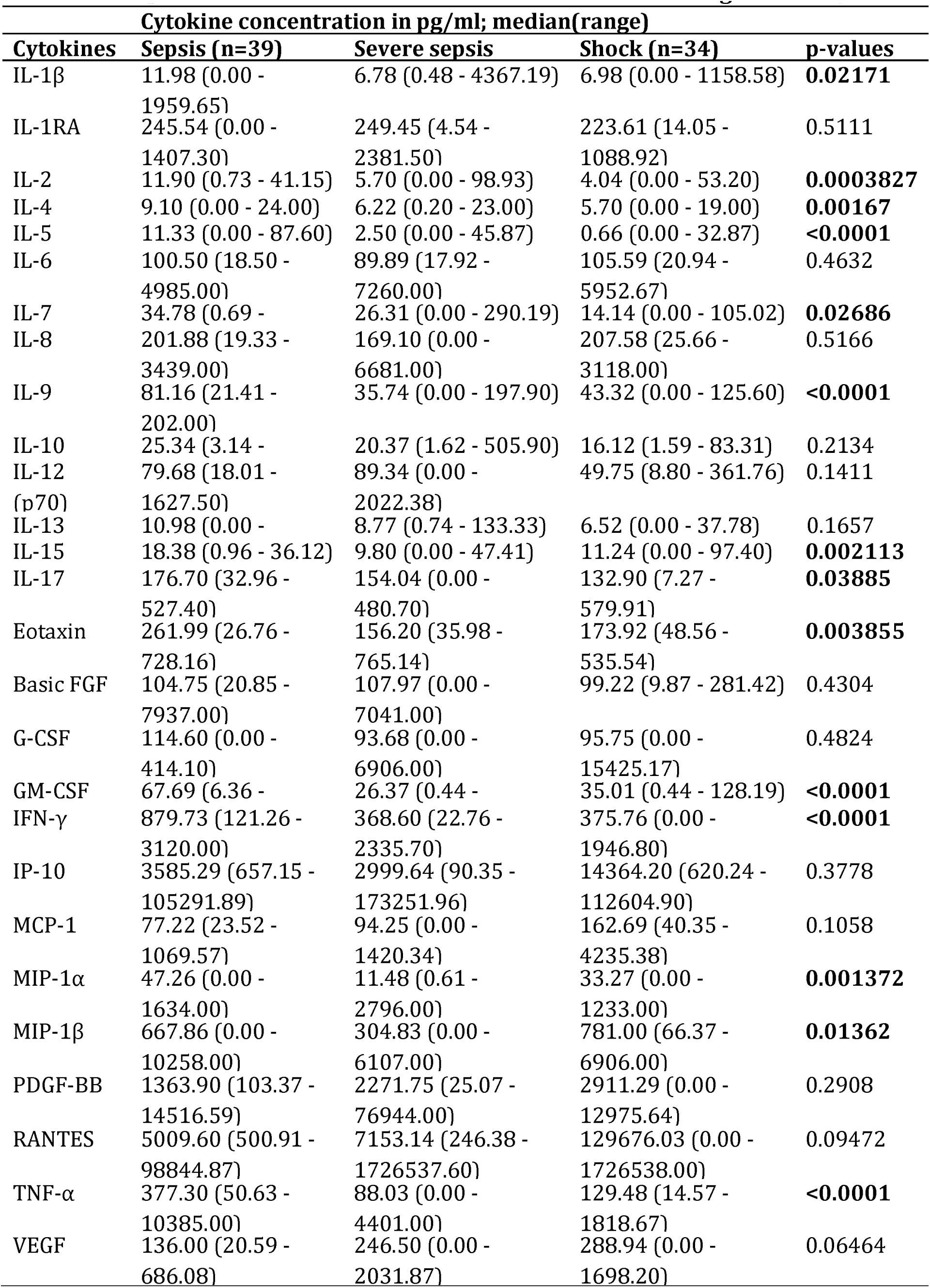
Comparison of plasma cytokine levels between clinical categories of sepsis

### Association of cytokine response with clinical severity

Although individual cytokines could discriminate between different forms of severity and outcome in human sepsis, more complex statistical modelling needed to be performed to extract valuable biological insight as cytokines usually exert their function as a group. Therefore, all plasma cytokine concentrations were loge (*n*+1) transformed and a 27X27 dimensional correlation matrix was constructed in order to examine associations between measured cytokines. The correlation matrix displayed in Figure 1 reveals existence of significant correlations between many of the 27 cytokines. This suggested that univariate analysis alone will be insufficient to extract all available information from the dataset. Therefore, to study effects of partitioning 139 cases into groups according to mortality or severity on combined host immune response, multivariate analysis of variance (MANOVA) was employed. The results of MANOVA on effects of the two main factors – outcome and clinical category – on observed variations among the twenty seven measured cytokines are depicted in Table 4. The results indicate that categorization based both on mortality as well as severity contributes significantly to differences in studied immune response variables. Moreover, there is also indication that categorization of the 139 sepsis cases into clinical stages of sepsis accounts for greater variation among the measured cytokines (approximately 51% as given by the value of Wilk’s lambda statistic that indicates proportion of variation not explained by a factor) as compared to categorization based on outcome viz., mortality (approximately 29%, Wilk’s lambda = 0.71525). Pair-wise contrasts between the three clinical categories to determine which of the individual categories differed from one another in terms of combined cytokine response are shown in Table 5 and revealed that in comparison to sepsis (category A), both severe sepsis (category B) and septic shock (category C) differed significantly in their combined immunological responses. The data also demonstrate that patients with severe sepsis or septic shock were comparable in their immune response. Similar results were obtained from a comparison of APACHE II scores between clinical categories of sepsis that showed significant differences between sepsis and severe sepsis or sepsis and septic shock but not between severe sepsis and septic shock (Figure 2).

**Table 4.** Multivariate analysis of the effects of outcome and severity on combined host immune response variables

| Source | Wilk's<br>lambda | F | Num. df | Den. Df | Pr>F |
| --- | --- | --- | --- | --- | --- |
| Outcome | 0.71525 | 1.6367 | 27 | 111 | **0.03950** |
| Category | 0.48783 | 1.7590 | 54 | 220 | **0.00246** |

**Table 5.**
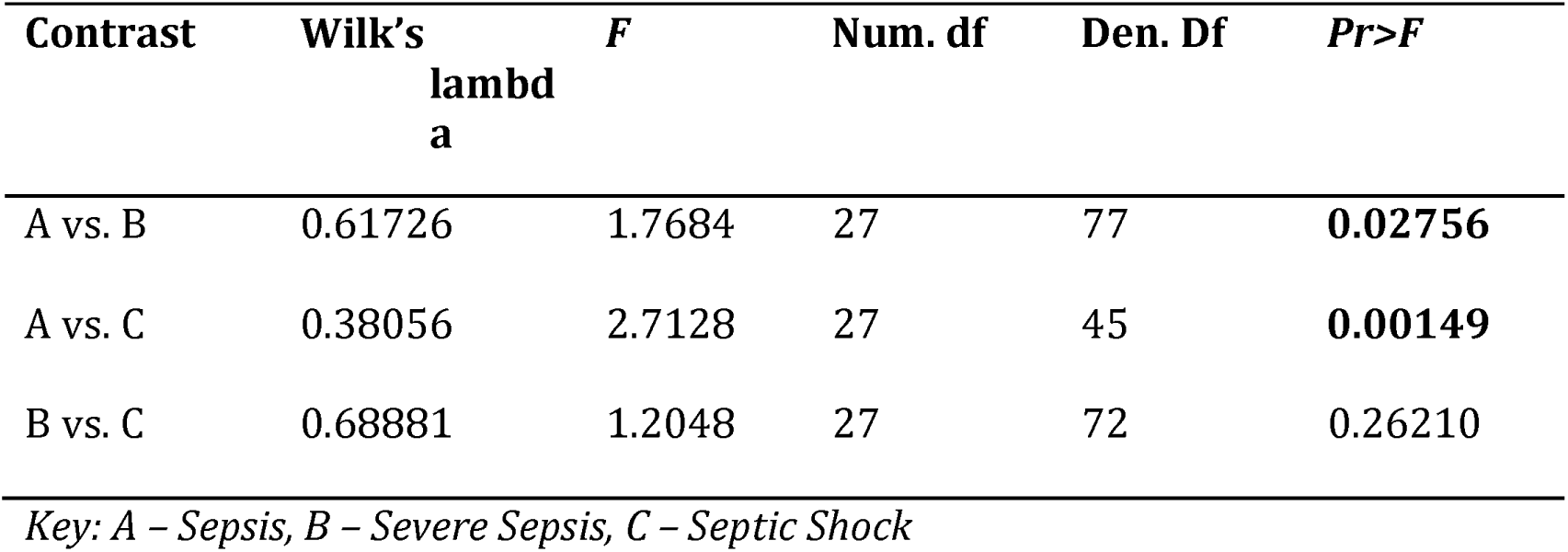
Pair-wise contrasts of three clinical categories of sepsis based on combined (multivariate) host immune response variables

| Contrast | Wilk's<br>lambda | F | Num. df | Den. Df | Pr>F |
| --- | --- | --- | --- | --- | --- |
| A vs. B | 0.61726 | 1.7684 | 27 | 77 | **0.02756** |
| A vs. C | 0.38056 | 2.7128 | 27 | 45 | **0.00149** |
| B vs. C | 0.68881 | 1.2048 | 27 | 72 | 0.26210 |

*Key: A – Sepsis, B – Severe Sepsis, C – Septic Shock*

**Figure 1.**
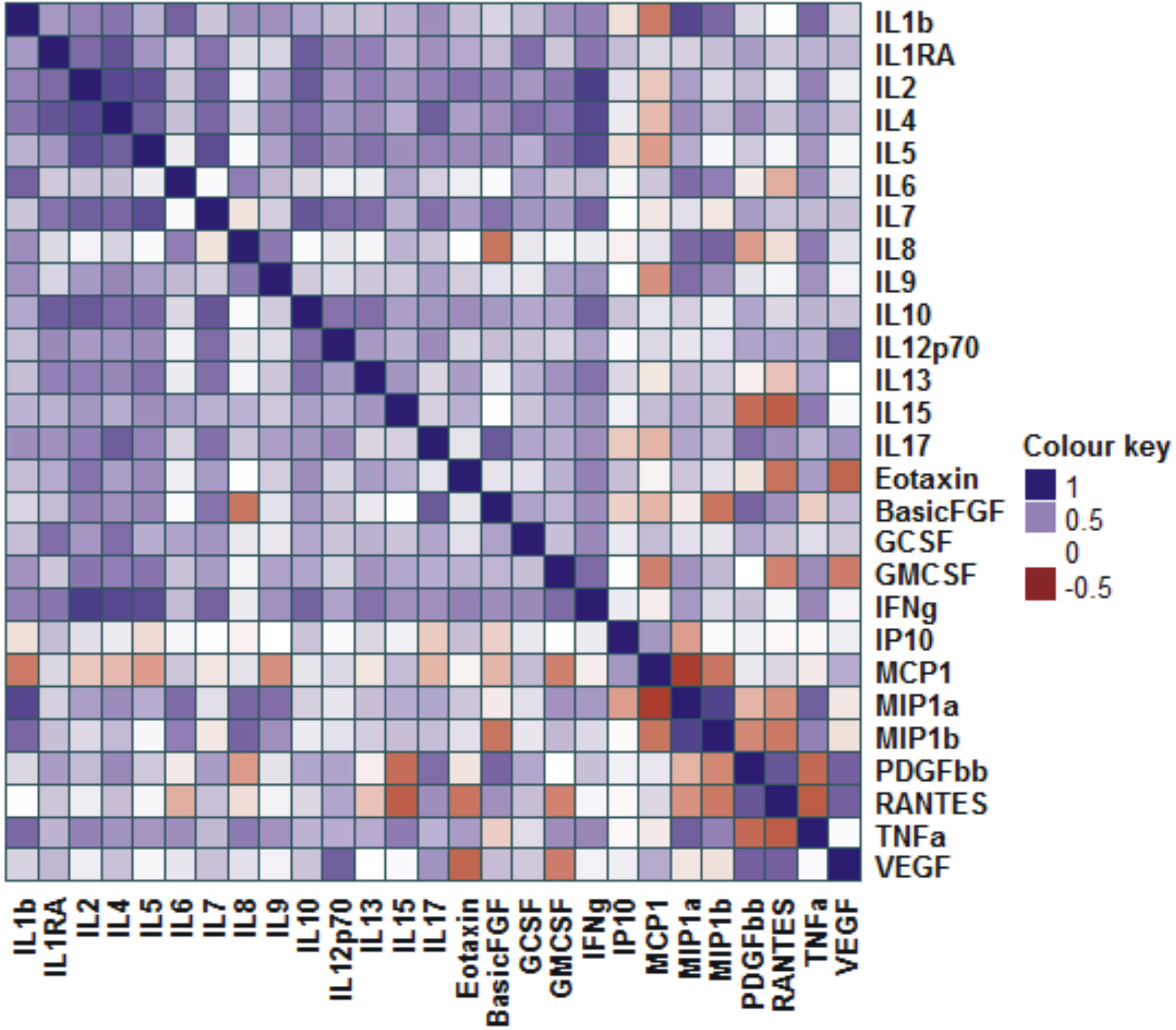
Associations between plasma cytokines in patients with sepsis. A 27 X 27 dimensional correlation matrix was constructed with loge (n+1) transformed values of plasma cytokines. The cytokines were clustered based on the correlation values. The colours represent the strength of the association. The correlation matrix was built in R statistical programming environment using corrplot and Heatmap packages.

**Figure 2.**
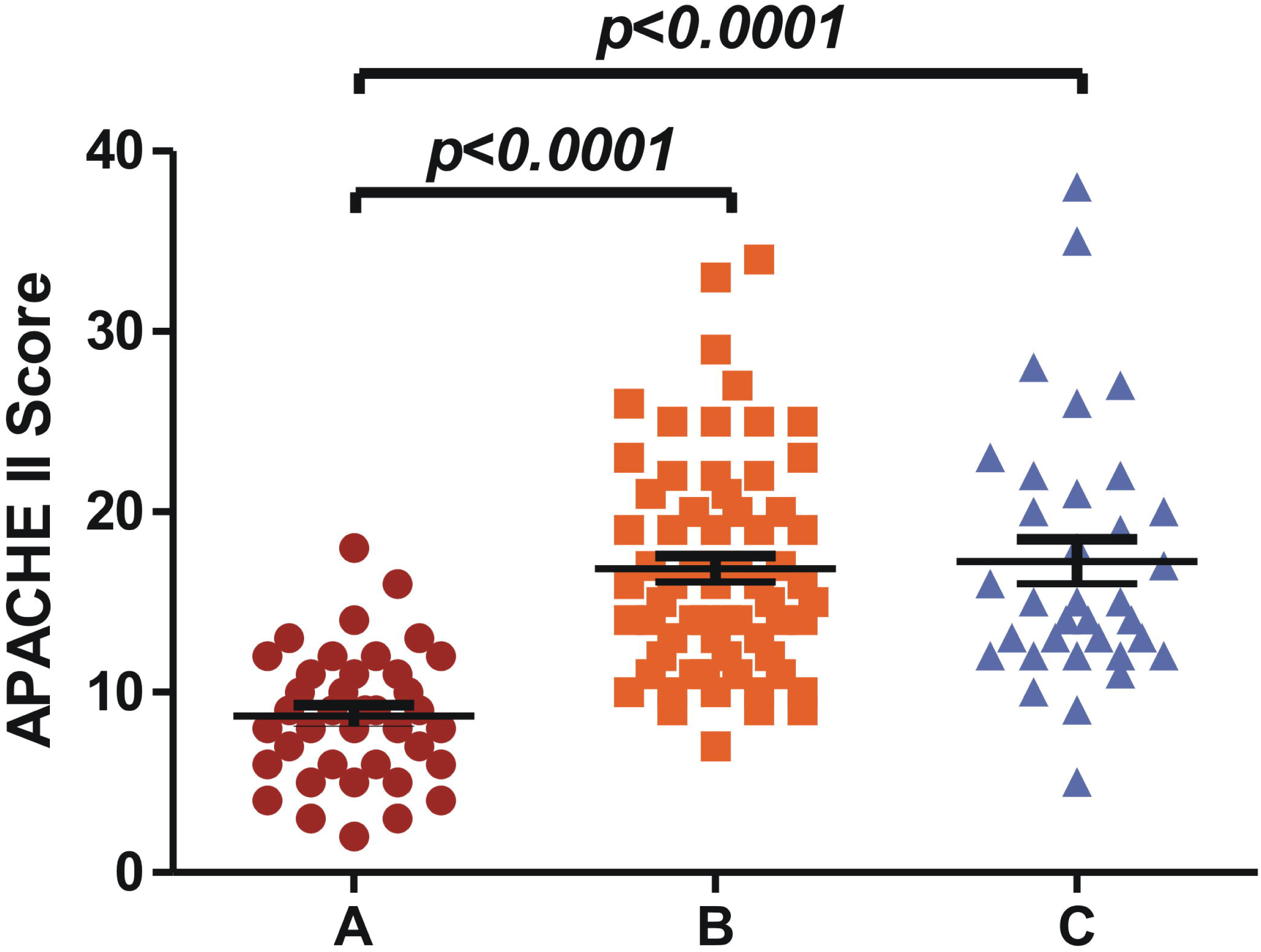
Comparison of APACHE II scores between clinical categories of sepsis. A: Sepsis (n = 39), B: Severe sepsis (n = 66) and C: Septic shock (n = 34). Significance was assessed using nonparametric one – way ANOVA (Kruskal – Wallis test) followed by Dunn’s multiple comparison to assess significance between individual groups.

### Immunological markers of disease severity and outcome

The results of canonical analysis to determine which combinations of the twenty seven cytokines are responsible for the estimated differences among the present outcome/severity groups are summarized in Tables 6 and 7. A canonical variate is a linear combination of dependent variables that can help discriminate between categories based on measured multiple response variables. Table 6 shows the two significant canonical variates derived for clinical severity. Can1, with an eigenvalue of 0.57 and accounting for 64.89% of explainable variation and about 0.36 of the total variation in the group of cytokines studied, may represent the composite variate that produces the greatest amount of among-category to within-category variation. The standardized canonical coefficients displayed in the table indicate the correlations between the two estimated variates and each of the individual cytokines examined. For canonical variate 1, these results show that TNF-α and IL-5 contribute most to among-category discrimination with respect to clinical severity. Moreover, the same sign for the standardized canonical coefficients for these two cytokines indicate a positive correlation. However, a similar canonical analysis conducted for outcome revealed an inverse association between IL-2 and IL-4, the cytokines that contributed most to among-category difference in terms of outcome (Table 7). This suggested a clear dichotomy of IL-2 and IL-4 in determining outcome in patients with sepsis. Figure 3 shows significantly higher IL-2:IL-4 ratio among survivors as compared to nonsurvivors, suggesting that higher plasma levels of IL-2 could be beneficial to the host.

**Table 6.** Canonical analysis showing the parameters and statistics of the significant canonical variates for sepsis immune response data based on clinical categories

| **Parameter** | **Can1** | **Can2** |
| --- | --- | --- |
| Eigen-value | 0.56780 | 0.30751 |
| Percent (explainable variation) | 64.868 | 35.132 |
| Squared canonical correlation (total variation | 0.36216 | 0.23519 |
| **Standardized canonical coefficients** |  |  |
| IL-1β | 0.106324 | 0.323111 |
| IL-1RA | 0.215646 | -0.507838 |
| IL-2 | 0.165034 | 0.272089 |
| IL-4 | 0.203382 | 0.065528 |
| IL-5 | **-0.586674** | -0.651274 |
| IL-6 | 0.358359 | 0.306899 |
| IL-7 | 0.162638 | -0.081257 |
| IL-8 | 0.265629 | -0.147053 |
| IL-9 | -0.265415 | 0.334220 |
| IL-10 | 0.311519 | -0.121265 |
| IL-12(p70) | -0.260803 | 0.076152 |
| IL-13 | 0.278206 | 0.251135 |
| IL-15 | -0.187366 | -0.141251 |
| IL-17 | -0.024788 | **1.000924** |
| Eotaxin | 0.103984 | 0.414003 |
| Basic FGF | -0.113082 | -0.103841 |
| G-CSF | -0.010419 | 0.005069 |
| GM-CSF | -0.012064 | 0.277336 |
| IFN-γ | -0.487986 | -0.135397 |
| IP-10 | -0.262748 | -0.015442 |
| MCP-1 | 0.037080 | 0.520671 |
| MIP-1α | 0.049497 | - |
| MIP-1β | -0.364949 | 0.665151 |
| PDGF-BB | -0.314133 | -0.362639 |
| RANTES | 0.079546 | -0.159529 |
| TNF-α | **-0.725480** | -0.125980 |
| VEGF | 0.385244 | -0.343296 |

**Table 7.** Canonical analysis showing the parameters and statistics of the significant canonical variate for sepsis immune response data based on outcome (survival vs mortality)

| Parameter | Can1 |
| --- | --- |
| Eigen-value | 0.39811 |
| Percent (explainable variation) | 100 |
| Squared canonical correlation (total variation explained) | 0.28475 |
| **Standardized canonical coefficients** |  |
| IL-1β | -0.2590933 |
| IL-1RA | 0.1856226 |
| IL-2 | - |
| IL-4 |  |
| IL-5 | -0.1021404 |
| IL-6 | 0.6371558 |
| IL-7 | 0.3968650 |
| IL-8 | -0.1665062 |
| IL-9 | 0.0363494 |
| IL-10 | 0.6812939 |
| IL-12(p70) | -0.4574752 |
| IL-13 | -0.2065759 |
| IL-15 | 0.1957445 |
| IL-17 | -0.3652665 |
| Eotaxin | 0.0565699 |
| Basic FGF | -0.0041714 |
| G-CSF | -0.6241971 |
| GM-CSF | 0.1069688 |
| IFN-γ | -0.2087967 |
| IP-10 | -0.5185169 |
| MCP-1 | 0.2661771 |
| MIP-1α | -0.0808227 |
| MIP-1β | -0.0562461 |
| PDGF-BB | -0.0437463 |
| RANTES | -0.2894390 |
| TNF-α | -0.1280161 |
| VEGF | -0.0211665 |

**Figure 3.**
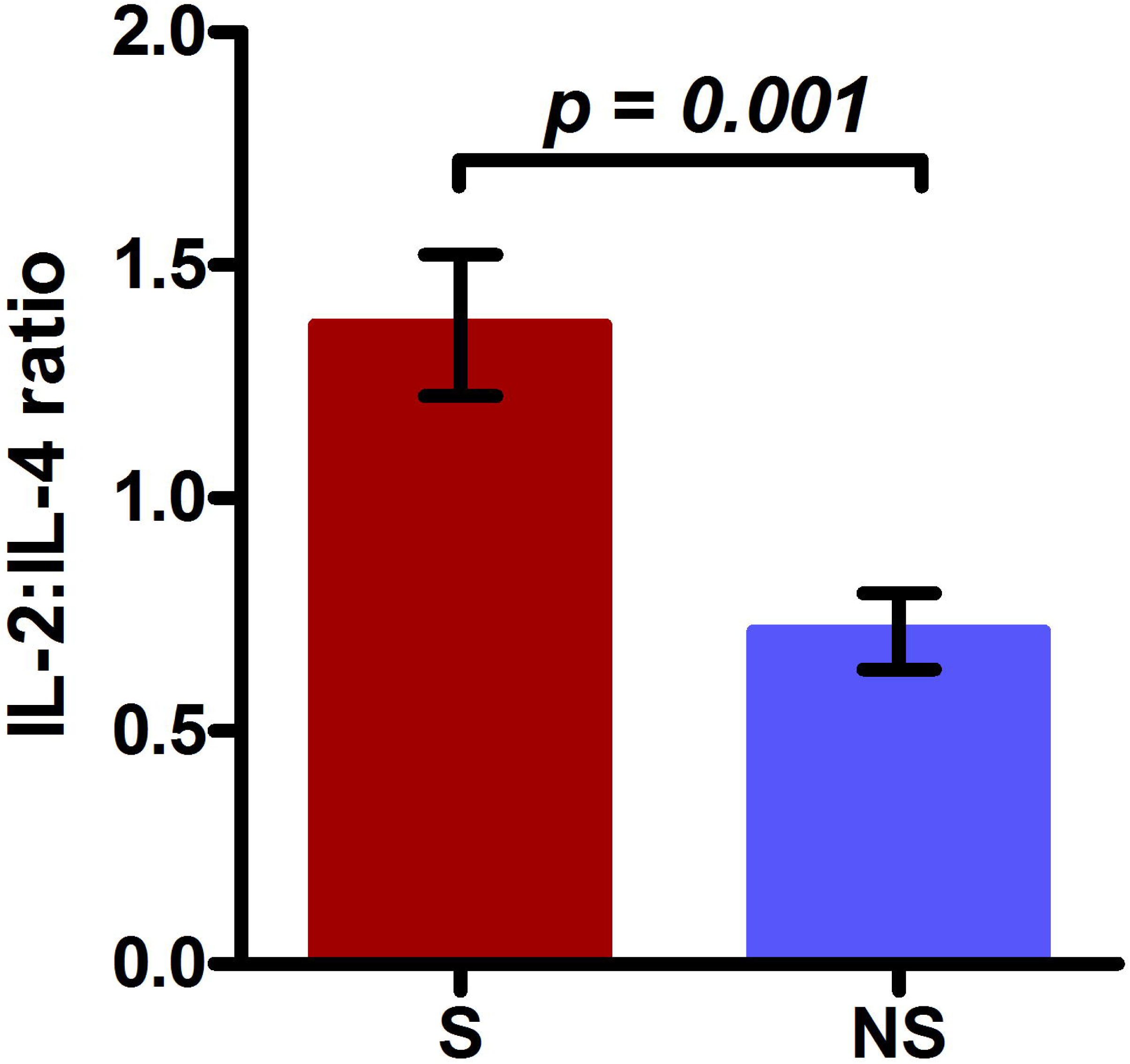
Comparison of IL-2:IL-4 ratio between survivors and nonsurvivors. S: Survivors (n = 91) and NS: Nonsurvivors (n = 48). Significance was assessed by unpaired t-test.

### Associations between different classes of immune response mediated by cytokines in human sepsis and mouse model of endotoxemia

To investigate interactions between different classes of host immune response and clinical pathology during sepsis, partial least squares – based structural equation model (PLS-SEM) was constructed. Structural equation modelling is a powerful statistical tool that enables modelling of complex multiple relationships among studied variables in a simplified path diagram. The results are shown in Figure 4. It is clear from the model that compensatory anti-inflammatory responses correlate more significantly with sepsis-associated clinical outcome (CARS 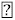 Clinical outcome, regression weight = 0.581, p-value = 0.029). Also, among the three cytokines contributing to the latent variable CARS, IL-4 had the highest factor loading that is suggestive of its strongest contribution to clinical outcome. Another interesting observation that came out from the model is a significant positive interaction between the latent variables SIRS (systemic inflammatory response syndrome) and CARS (compensatory anti-inflammatory response syndrome; regression weight = 0.609, p-value <0.0001). This suggested simultaneous regulation of inflammatory and anti-inflammatory components of the host response in sepsis. Further validation for concurrent activation of pro- and anti-inflammatory cytokine response in sepsis was obtained by induction of inflammation in C57BL/6 mice through injection of lipopolysaccharide (LPS) and measuring of cytokines in plasma. Two doses of LPS – 5 mg/kg (nonlethal dose) and 35 mg/kg (lethal dose) – were used to simulate cases of survivors and nonsurvivors in human sepsis. Data presented in Figure 5 reveals that irrespective of dose of LPS, pro- and anti-inflammatory cytokines displayed similar kinetic profiles by peaking at 2 – 4 hours post injection followed by a gradual decline. PLS-SEM analysis also confirmed the strong positive association between inflammatory and immunosuppressive mediators (Figure 6, SIRS 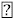 CARS, regression weight = 0.862, p-value <0.0001). Taken together, above data suggest that pro- and anti – inflammatory response during human sepsis and murine endotoxemia are stimulated and regulated simultaneously with anti – inflammatory cytokine levels contributing more significantly towards severe clinical outcome in human sepsis.

**Figure 4.**
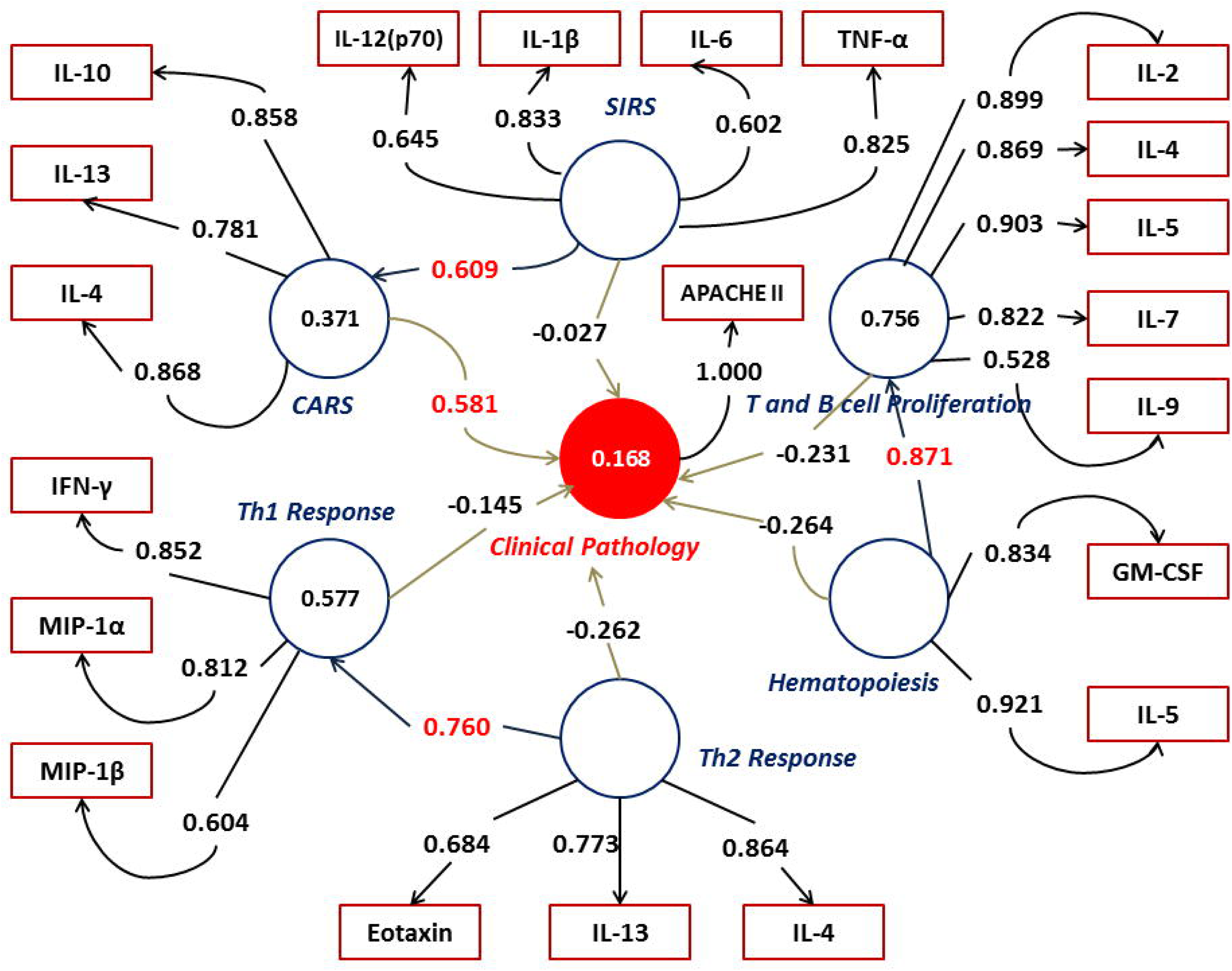
Partial least squares regression – based Structural equation modelling (PLS-SEM) of immune response data in patients with sepsis. The complex inter-relationships amongst the components of immune response to sepsis in humans were analysed by structural equation modelling. Circles represent latent variables and rectangles represent measured variables. Numbers inside the circles denote R^2^ values that indicate variance of each factor. The strength of interaction between two latent variables is given by the regression weight. Numbers associated with measured variables are the factor loadings for each variable which is a reflection of each measured variable’s contribution to the observed variance.

**Figure 5.**
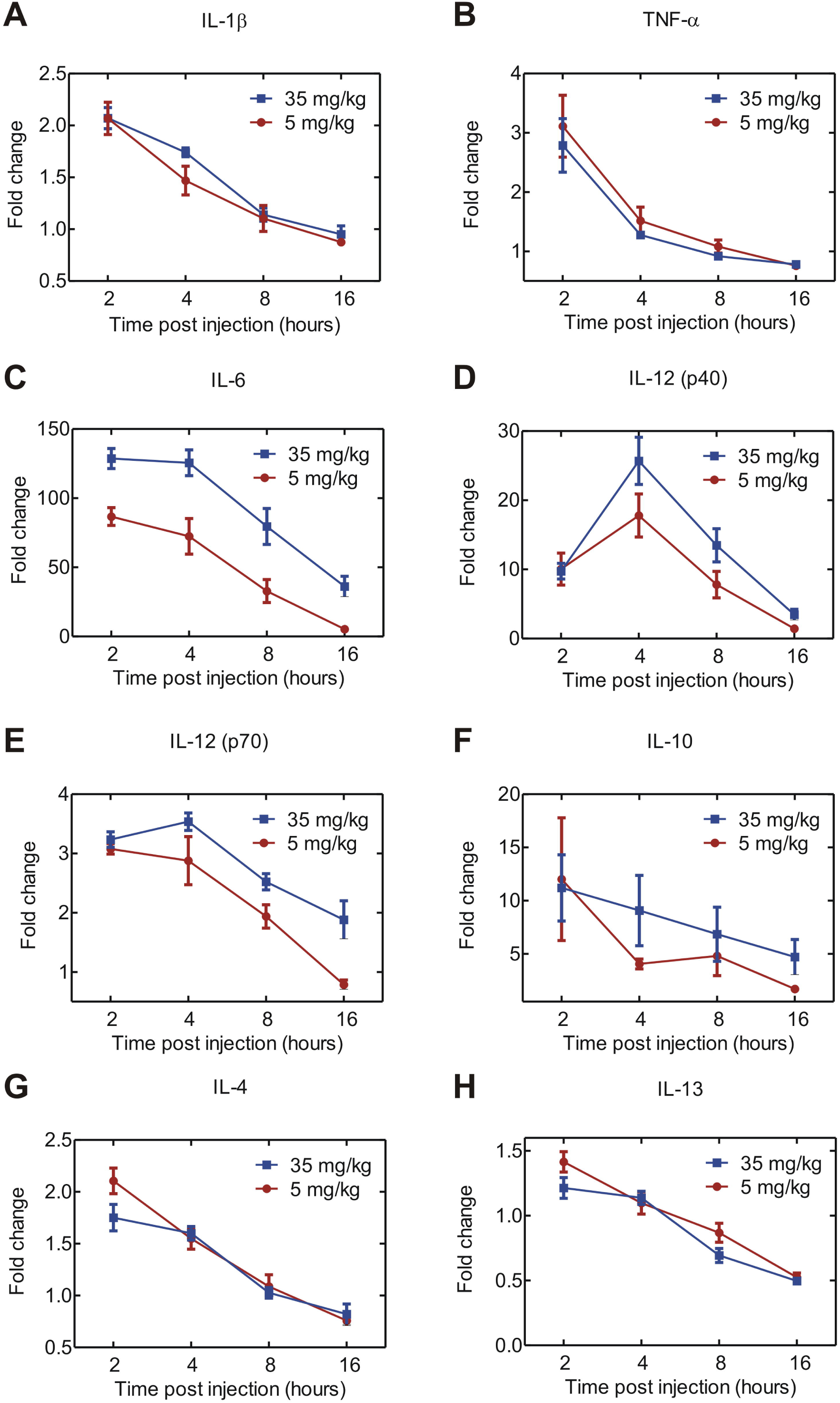
Temporal profile of plasma levels of key inflammatory (A-E) and anti-inflammatory (F-H) cytokines in C57BL/6 mice based on LPS dose. Different batches of mice were injected with two different doses of LPS as indicated. The animals were sacrificed at the indicated time points and cytokines in plasma were measured by a Bioplex mouse 23 – plex kit (Bio – Rad). Data is shown as fold change over uninjected control mice. Each data point is mean ± S.E.M. of seven animals.

**Figure 6.**
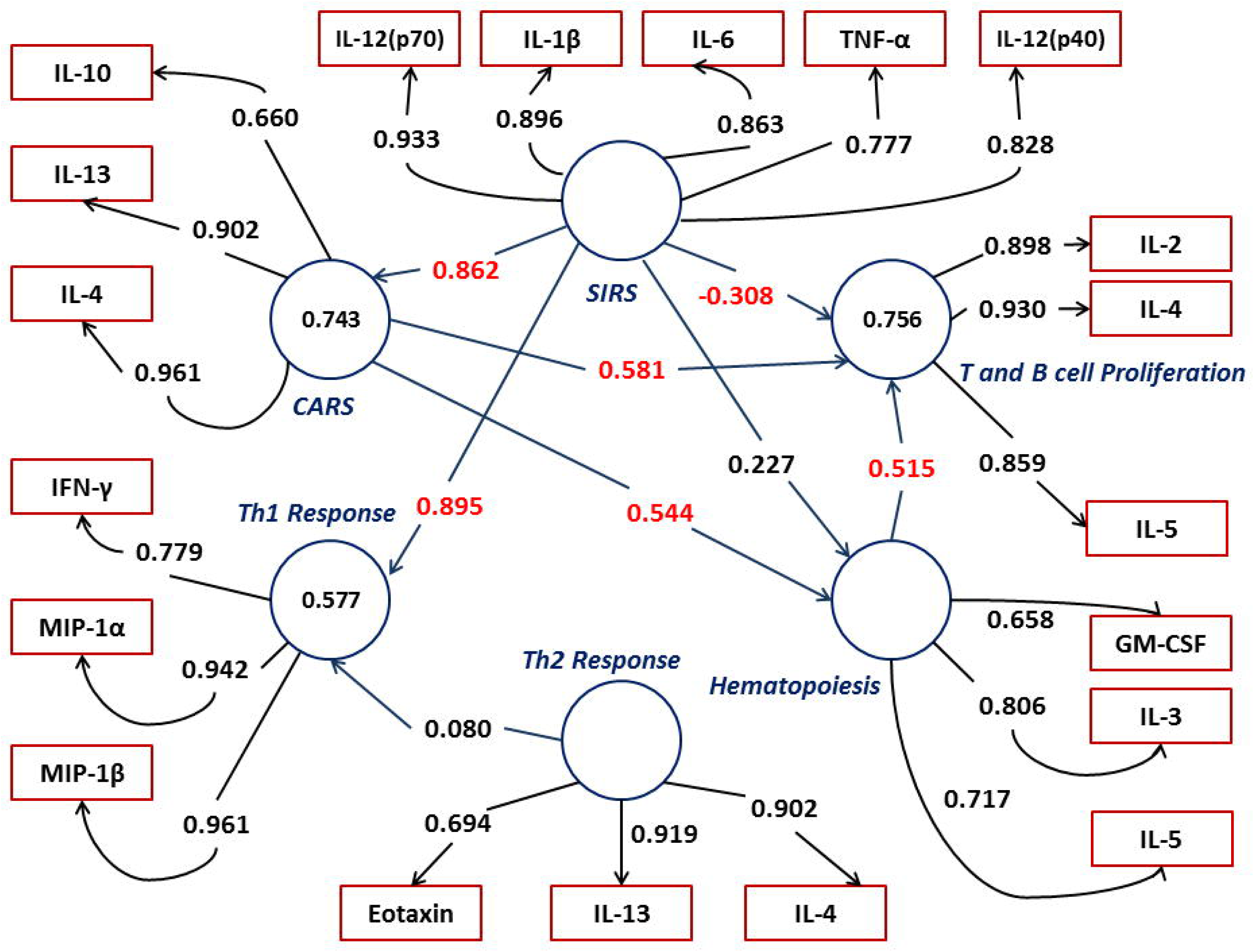
Partial least squares regression – based Structural equation modelling (PLS-SEM) of immune response data in endotoxic mice. The complex inter-relationships amongst the components of immune response in mice to LPS-induced endotoxemia were analysed by structural equation modelling. Circles represent latent variables and rectangles represent measured variables. Numbers inside the circles denote R^2^ values that indicate variance of each factor. The strength of interaction between two latent variables is given by the regression weight. Numbers associated with measured variables are the factor loadings for each variable which is a reflection of each measured variable’s contribution to the observed variance.

## Discussion

The first significant observation from the present study is the ability of IL-1β, IL-2, IL-4, IL-5, IL-17 and IFN-γ to discriminate survivors from nonsurvivors and also between clinical categories of sepsis. Additionally, plasma levels of BasicFGF and RANTES could segregate survivors from nonsurvivors whereas IL-7, IL-9, IL-15, Eotaxin, GM-CSF, MIP-1α, MIP-1β and TNF-α levels were successful in contrasting different clinical stages of sepsis. Owing to strong collinearity between many of the measured cytokines, multivariate statistical methods were used for further analyses. Multivariate analysis of variance (MANOVA) conducted with plasma cytokine values as dependent variables revealed that stratification of patients based on clinical severity had a greater impact on observed differences between cytokine levels as compared to partitioning by outcome. Pair-wise comparison of multiple cytokine response between the three clinical categories revealed a clear distinction between sepsis and severe sepsis or septic shock. However, immune response mediated by cytokines in plasma was comparable between severe sepsis and septic shock. The same was true with respect to clinical score as well, as APACHE II scores could only discriminate between sepsis and severe sepsis or septic shock and not between severe sepsis and septic shock. Thus, it appears that clinically and immunologically, severe sepsis and septic shock are virtually indistinguishable. In fact, the window of time in which a patient can move from severe sepsis to shock could be very narrow. Although it is appealing to device biomarkers that can differentiate these two late stages of sepsis, however, given the high mortality rates associated with both the stages, it is probably safe to assume that such a marker would probably be of very little practical use. Thus, it appears that clinically and immunologically, severe sepsis and septic shock are virtually indistinguishable. In fact, the window of time in which a patient can move from severe sepsis to shock could be very narrow. Although it is appealing to device biomarkers that can differentiate these two late stages of sepsis, however, given the high mortality rates associated with both the stages, it is probably safe to assume that such a marker would probably be of very little practical use. These observations are in conformity to the recent clincial classification of Sepsis by a consortium of investigators recommonding that Severe Sepsis and Shock should be merged as a single category (14a)

A key finding of the current study came through the use of canonical analysis to investigate the relative importance of measured cytokines responsible for the estimated differences between the study groups based on MANOVA results. It was observed that IL-2 and IL-4 were inversely associated in the canonical analysis. Therefore, comparison of IL-2:IL-4 ratio between survivors and nonsurvivors revealed lower IL-2:IL-4 ratio to be associated with bad prognosis. Thus, it can be concluded that higher IL-2 is associated with better prognosis, a finding in contrast with previously published report that showed increased IL-2 in parallel with disease severity [21]. Concurrently, it is also evident that higher IL-4 is detrimental to the host, an observation in accordance with an earlier report [22].

Modulation of inflammatory signalling during sepsis has generally been considered as a possible intervention strategy to improve survival in sepsis and prevent septic shock. Even though results of anti-inflammatory interventions in animal models [23-25] and human phase I and II trials [26, 27] were promising, phase III clinical trials failed demonstrable success [28-30] and in some cases such as iNOS inhibitors, were even detrimental [31]. These interventions were based on the logic that immune response in sepsis represents an interplay of two contrasting phenomena related to the inflammatory status of the patient. The early systemic inflammatory response syndrome (SIRS) is characterized by overproduction of proinflammatory cytokines (hyperinflammatory status), which then is progressively suppressed by development of an compensatory anti-inflammatory response (hypoinflammatory status) syndrome (CARS) [10, 32]. This hypothesis was further enriched by introduction of the expression ‘mixed anti-inflammatory response syndrome’ (MARS) that represents temporary homeostasis between decreasing SIRS and increasing CARS [10, 33]. Although this multimodal nature of immune response during sepsis is a widely accepted paradigm currently, there is no experimental evidence to convincingly prove it. On the contrary, in recent years, work based both in animal models and in human sepsis have suggested simultaneous progression of pro- and anti-inflammatory responses [34, 35]. In the present study, structural equation modelling was used to assess the various interactions between different facets of immune response in sepsis patients and their impact on clinical pathology. Such an analysis has not been reported in literature. Structural equation modelling is a powerful statistical tool that combines statistical modelling with path diagram to test hypotheses consisting of interacting variables and pathways with reference to available theoretical knowledge [36-38]. In a biological system, there are complex interactions between causes and effects. For instance, the effects of one factor may be explained by another factor directly as well as indirectly by a third factor via the effect of a fourth factor. Furthermore, the first factor itself may be an explanatory variable for other factors. These complex inter-relationships cannot be fully explored by standard correlation or regression analysis. Structural equation modelling offers the advantage of dissecting these relationships in a path diagram by a system biology approach thus assessing the total effects of variables on one another.

Using a partial least squares regression – based algorithm to fit the data, the constructed structural equation model revealed two important findings. Firstly, a strong positive association was observed between the latent variable CARS consisting of anti-inflammatory cytokines as measured component variables and the latent variable clinical pathology, which had APACHE II scores as it components. This strongly suggests a direct effect of anti-inflammatory cytokines as chief contributors of clinical pathology in human sepsis. Secondly, strong positive association was observed between inflammatory and anti-inflammatory cytokines that suggested simultaneous rather than sequential regulation of these two contrasting arms of the immune system in sepsis. This observation in human sepsis cases was further tested in a mice model of acute inflammation that shows a similar concurrent regulation of inflammatory and anti-inflammatory responses during the course of inflammation. Two earlier reports have also demonstrated similar observations in mice models of polymicrobial sepsis induced either by cecal ligation and puncture (CLP) or colon ascendens stent peritonitis (CASP) [35, 39]. Observations reported from our laboratory earlier also offers further support. When normal human blood was stimulated in vitro with LPS and intracellular cytokines were quantified using multicolour flowcytometry both inflammatory and anti-inflammatory cytokines (TNF, IL1-b and IL-10) were simulataneusly stimulated [40]. The results of the current study suggesting that host molecules associated with CARS are more critical for bad prognosis in human sepsis is not surprising in the context of the following reasons: (a) results of several clinical trials directed towards use of drugs/antagonists of inflammatory host response have consistently failed [8, 41]; (b) increased proinflammatory cytokines are very often associated with increase in anti-inflammatory cytokines such as IL-10 [42]; (c) high TNF-α:IL-10 ratio correlated better with increased mortality [42]; and (d) in vitro activation of immune cells from sepsis patients by endotoxin led to release of very low levels of inflammatory cytokines such as TNF-α, IL-1β and IL-6 in comparison to healthy individuals [43]. These findlings have immediate clinical implication for treatment of sepsis. As per the model presented in this study, treatment of patients targeting one of the immune mechanisms (only inflammatory or only antiinflamamtory) may be counterproductive. One possible course of action, therefore, could be combination therapeutic strategy targeting either pro- and anti-inflammatory mechanisms after careful analysis of the status of inflammation parameters in each of the patients, for eg., definition of sepsis cases based on decreased HLA-DR and increased PD-1R expression indicative of immunosuppressed state could be used a biomarker to use immuno-stimulatory molecules.

## Conclusions

In the present study, we provide empirical evidence in support of concurrent induction of pro and anti – inflammatory cytoine responses in human sepsis and experimental mouse endotoxemia. Using robust statistical modelling, the study also predicts a critical role for anti – inflammatory cytokines in the pathogenesis of human sepsis. Despite multiple efforts from several research groups over a long period of time, sepsis continues to be a major global health burden. The past failures of therapeutic strategies based on targeted inhibition of inflammation called for a better understanding of the host response in sepsis. As such, the conclusions drawn from the current study should offer novel insights into the pathogenesis of human sepsis that could lead to design of improved therapeutic strategies.

## Abbreviations

SIRS: systemic inflammatory response syndrome
CARS: compensatory anti – inflammatory response syndrome
MARS: mixed antagonist response syndrome
CLP: cecal ligation and puncture
CASP: colon ascendens stent peritonitis
APACHE: acute physiology and chronic health evaluation
MANOVA: multivariate analysis of variance
PLS-SEM: partial least squares structural equation modelling.

## Ethics approval

Signed informed consent was obtained from all participants. The study was approved by ethics committees of Institute of Life Sciences, Bhubaneswar and S.C.B. Medical College, Cuttack.

All animal experiments were approved by the institutional animal ethics committee of Institute of Life Sciences, Bhubaneswar.

## Availability of data and materials

The datasets upon which the conclusions of this article are based are available as supplementary files accompanying this manuscript.

## Competing interests

The authors declare no competing financial interests.

## Funding

The Institute of Life Sciences is funded by grants from Department of Biotechnology, Govt. of India. BR was funded by DBT, Ministry of Science and Technology for a project titled “Reprogramming Deleterious Inflammatory Monocytes in Sepsis”. RM is a recipient of a Senior Research Fellowship from the Indian Council of Medical Research, Govt. of India.

## Author Contributions

RM executed most of the experiments, performed statistical modelling, and prepared the draft manuscript; PKB and BKP analysed plasma levels of cytokines; PKT and BKD performed all clinical evaluation and categorization of patients; RT conducted biochemical assays and laboratory diagnostic tests; and BR conceived the project, interpreted data and finalized the manuscript.

## Additional files

**Additional file 1**: BMC_Inf Dis _Hu_Dataset.xlsx. This file contains the plasma cytokine concentrations measured in 139 human sepsis patients.

**Additional file 2**: BMC_Inf Dis_Mo_Dataset.xlsx. This file contains the plasma cytokine concentrations measured in C57BL/6 mice injected with two different doses of LPS.

**Additional file 3**: BMC_Inf Dis_R-scripts.txt. This file contains the R commands used for the statistical analyses performed in the manuscript.

